# Towards Efficient *k-*Mer Set Operations via Function-Assigned Masked Superstrings

**DOI:** 10.1101/2024.03.06.583483

**Authors:** Ondřej Sladký, Pavel Veselý, Karel Břinda

## Abstract

The design of efficient dynamic data structures for large *k*-mer sets belongs to central challenges of sequence bioinformatics. Recent advances in compact *k*-mer set representations via Spectrum-Preserving String Sets (SPSS), culminating with the masked superstring framework, have provided data structures of remarkable space efficiency for wide ranges of *k*-mer sets. However, the possibility to perform set operations with the resulting indexes has remained limited due to the static nature of the underlying compact representations. Here, we develop *f*-masked superstrings, a concept combining masked superstrings with custom demasking functions *f* to enable *k*-mer set operations based on index merging. Combined with the FMSI index for masked superstrings, we obtain a memory-efficient *k*-mer membership index and compressed dictionary supporting set operations via Burrows-Wheeler Transform merging. The framework provides a promising theoretical solution to a pressing bioinformatics problem and highlights the potential of *f*-masked superstrings to become an elementary data type for *k*-mer sets.

## 1 Introduction

To store and analyze the vast volumes of DNA sequencing data [41], modern bioinformatics methods increasingly rely on *k-mers, k*-long substrings of genomic data. As *k*-mer-based methods bypass the computationally expensive sequence alignment, they become increasingly popular in data-intense applications such as large-scale data search [5,13,23], metagenomic classification [40,11], infectious disease diagnostics [6,12], or transcript abundance quantification [7,30]. All these applications rely on indexing collections of *k*-mer sets using advanced data structures [26], which often rely at their core on data structures for single *k*-mer sets [15].

A central challenge in contemporary sequence bioinformatics is to design single-*k*-mer-set data structures with two qualities: scalability and dynamism. On one hand, as *k*-mer sets can be large, possibly exceeding billions of distinct *k*-mers [23], we need data structures that are efficient both in space and time simultaneously, with an internal adaptability to different types of *k*-mer sets. On the other hand, as modern genomic databases undergo rapid development, due to rapidly growing content as well as databases curation, we also need to perform efficient updates across *k*-mer indexes to avoid repetitive and costly index recomputations. This includes scenarios such as *k*-mer set operations across sets, and additions or removals of individual *k*-mers.

A substantial advance in the scalability challenge was achieved using the concept of Spectrum-Preserving String Sets (SPSS) [16,9,10,32,35,36]. *k*-mers in genomic *k*-mer sets tend to be highly non-independent [14] – they typically correspond to sets of *k*-long substrings of a small number of (potentially long) strings, a property known as the *spectrum-like property (SLP)* [15]. This gave rise to textual *k*-mer set representations that correspond to path covers of de Bruijn graphs, which we will collectively refer to as SPSS [16,9,10,32,35,36] and which are now standard and widely used across data structures (e.g., in [27,2,31]).

*Masked superstrings* (MS) provided additional space gains and structural adaptability by virtue of a better *k*-mer set compaction [37]. The core improvement over SPSS lies in modeling the structure of *k*-mer sets by overlap graphs instead of de Bruijn graphs, thus being able to exploit overlaps of *any* length. MS represent *k*-mer sets using an approximately shortest superstring of all *k*-mers and a binary mask to avoid false positive *k*-mers. MS generalize any existing SPSS representation as these can always be encoded as MS, but provide further compression power, especially for non-SLP data arising from sketching or subsampling. The resulting representation is well indexable using a technique called Masked Burrows Wheeler Transform [39], resulting in a *k*-mer data structure with 2 + *o*(1) bits per *k*-mer under the SLP.

However, the lack of dynamism of both SPSS and MS – and of their derived data structures – has been limiting their wider applicability. To the best of our knowledge, the only supported operations were union, either via merging SPSS of several *k*-mer sets resulting in an SPSS of their union, or by the Cdbg-Tricks [19] to calculate the union unitigs from the unitigs of multiple original *k*-mer sets. However, other operations besides union, such as intersection or symmetric difference, have never been considered.

Here, we develop a dynamic variant of masked superstrings (and thus also of SPSS) called the *f-masked superstrings (f-MS)*. The key idea is to equip masked superstrings with so-called demasking functions *f* for more flexible mask interpretation (**Sec. 3**). When complemented with the concatenation and mask recast operations (**Sec. 4.1**) and possibly compaction (**Sec. 4.2**), this provides support for union, intersection, and symmetric difference and thus any set operation with *k*-mer sets, resulting in a complete algebraic type for *k*-mer sets (**Sec. 4.3**). We demonstrate the applicability of the concept on the FMSI index for masked superstrings [39] (**Sec. 5**) and provide a proof-of-concept implementation in the FMSI program (**Sec. 6**).

### 1.1 Problem formulation

We focus on the problem of developing a time- and space-efficient index for *k*-mer sets with support for set operations.

i. **Index construction**, with time complexity linear in the number of *k*-mers.
ii. **Membership queries**, with low bits-per-*k*-mer memory requirements, approaching 2 bits per distinct *k*-mer for datasets following the SLP [15].
iii. **Set operations**, including union, intersection, difference, and symmetric difference.

### 1.2 Related Work

Many works have recently focused on data structures for single *k*-mer sets and their collections; we refer to [15,26] for recent surveys. Here, we primarily focus on those that are exact and offer some kind of dynamicity, i.e., an efficient support for set operations or, at least, insertions/deletions of individual *k*-mers. The recently introduced Conway-Bromage-Lyndon (CBL) structure [28] builds on the work of Conway and Bromage [17] on sparse bit-vector encodings and combines them with smallest cyclic rotations of *k*-mers (a.k.a. Lyndon words), which yields a dynamic and exact *k*-mer index supporting set operations, such as union, intersection, and difference, as well as insertions or deletions of *k*-mers.

While, to the best of our knowledge, other *k*-mer indexes do not support efficient set operations such as the intersection or difference, other tools, including BufBoss [1], DynamicBoss [3], and FDBG [18] allow for efficient insertions and deletions of individual *k*-mers. Bifrost [20], VARI-merge [29], Metagraph [23], or the very recent Cdbg-Tricks [19] support insertions but not deletions. We note that there are many more highly efficient but static data structures for individual or multiple *k*-mer sets, e.g., [4,25,2]. In particular, one can combine SPSS representations with efficient full-text search [16,32,10] or hashing [31], but this has so far yielded only static indexes.

Another line of work focused on *k*-mer counting, where we additionally require to compute the *k*-mer frequencies. Although some counters [22,24,33] support operations such as union, intersection, or difference on *k*-mer multisets, counting typically requires comparatively larger memory and heavy disk usage compared to efficient *k*-mer indexes.

#### Baseline approaches for performing set operations

We note that one can add support for set operations to a static data structure in a straightforward way: One option is extracting the *k*-mer sets from the input indexes, performing the given operation with the sets, and computing the new index for the resulting set; however, this process requires substantial time and memory. Another option is to keep the indexes for input *k*-mer sets and process a *k*-mer query on the set resulting from the operation by asking each index for the presence/absence of the *k*-mer in each input set, which is, however, time and memory inefficient as all the indexes need to be loaded into memory, as also noted in [21]. Therefore, we seek to perform set operations without the costly operation of recomputing the index or making multiple queries to original indexes.

## 2 Preliminaries

### Strings and *k*-mers

We use constant-size alphabets *Σ*, typically the nucleotide alphabet *Σ* = {`A`, `C`, `G`, `T`} (unless stated otherwise). The set *Σ*^*^ contains all finite strings over *Σ*, with *ε* representing the empty string. A substring of *S* is a contiguous sequence of characters within *S*. For a given string *S* ∈*Σ*^*^, |*S*| denotes its length, and |*S*| _*c*_ the number of occurrences of the letter *c* in *S*. For two strings *S* and *T, S* + *T* denotes their concatenation. A *k-mer* is a *k*-long string over *Σ*. For a string *S* and a fixed length *k*, the *k*-mers *generated* by *S* are all *k*-long substrings of *S*, and similarly for a set of strings *ℛ*, they are those generated by the individual strings *S*∈*ℛ* . We present our framework in the uni-directional model, where all *k*-mers are treated as distinct (unless stated otherwise).

#### Masked superstrings

Given a *k*-mer set *K*, a *masked superstring* (MS) [37] consists of a pair (*S, M* ), where *S* is an arbitrary superstring of the *k*-mers in *K* and *M* is a binary mask of the same length. An occurrence of a *k*-mer in an MS is said to be on if there is `1` at the corresponding position in the mask (i.e., the initial position of the occurrence), and off otherwise. The set of *k*-mers generated by *S* are referred to as the *appearing k-mers*, and only those that have at least one on occurrence are *represented k-mers*, i.e. those from the represented set *K*. All masks *M* that represent a given *K* in a combination with a given superstring are called *compatible*.

#### Occurrence function

Let (*S, M* ) be an MS and suppose that our objective is to determine whether a given *k*-mer *Q* is among the MS-represented *k*-mers. Conceptually, this process consists of two steps: (1) identify the starting positions of occurrences of *Q* in *S*, and (2) verify using the mask *M* whether at least one occurrence of *Q* is on. We can formalize this process via a so-called *occurrence function*.

##### Definition 1.

*For a superstring S, a mask M, and a k-mer Q, the* occurrence function *Λ*(*S, M, Q*) → { 0, 1 }^*^ *is a function returning a finite binary sequence with the mask symbols of the corresponding occurrences, i*.*e*.,

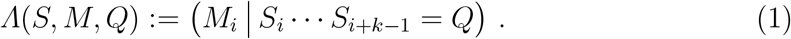

In this notation, verifying *k*-mer presence corresponds to evaluating the composite function ‘**or** ◦ *Λ*’; that is, a *k*-mer is present if *Λ*(*S, M, Q*) is non-empty and the logical **or** operation on these values yields 1. Thus, the set of all MS-represented *k*-mers can be expressed as

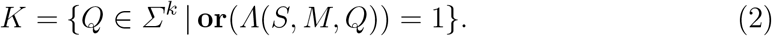

##### Example 2.

Consider the *k*-mer set *K* = { `ACG`, `GGG` }. One possible superstring is `ACGGGG`, with three compatible masks: `101100`, `100100`, `101000`, resulting in the mask-cased encodings `AcGGgg`, `AcgGgg`, `AcGggg`, respectively. Given a *k*-mer `GGG` and masked superstring `AcgGgg`, *Λ*(*S, M, Q*) = (0, 1) and **or**((0, 1)) = 1, therefore `GGG` is considered represented.

## 3 Function-Assigned Masked Superstrings

This work is based on two important observations on masked superstrings in the context of their applications for *k*-mer-set indexing.

First, **or** is one member of a large class of functions that could be used to demask *k*-mers in masked superstrings: for instance, MS could have been defined using the **xor** function, with a *k*-mer considered present if the number of its on occurrences is odd. In fact, any symmetric Boolean function *k*-mer demasking can serve the role.

Second, different data structures may impose different constrains on *f*-masked superstrings, and this can be embedded directly into demasking functions via a special return value, `invalid`. Additionally, we treat the non-appearing *k*-mers as non-represented by requiring *f* (*ε*) = 0.

### Definition 3.

*We call a symmetric function f* : {0, 1} ^*^→ {0, 1, `invalid`} *with f* (*ε*) = 0 *a k*-mer demasking function.

We now generalize the concept of MS into so-called *f*-masked superstrings (*f*-MS).

### Definition 4.

*Given a demasking function f, a superstring S, and a binary mask M, such that* |*M*| = |*S*|, *we call a triplet 𝒮* = (*f, S, M* ) *a* function-assigned masked superstring *or f*-masked superstring, *and abbreviate it as f*-MS.

*If f* (*Λ*(*S, M, Q*)) = `invalid` *for any k-mer Q, we call the f-MS* invalid.

Now, for a valid *f*-MS, we generalize Equation (2) for *k*-mer decoding as

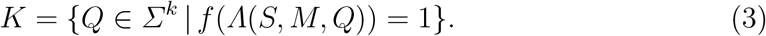

We note that by the validity requirement, *f* imposes structural constraints on the valid masks, e.g., **oon** requires that only a single occurrence of a represented *k*-mer is on and all other occurrences are off. Still, there may be many valid masks, leaving room for an application-specific mask optimization. In Table 1 we overview *f*-MS used throughout the paper. See also Figure 1a for an example evaluation of *k*-mer presence for sample *f*-MS.

**Table 1:**
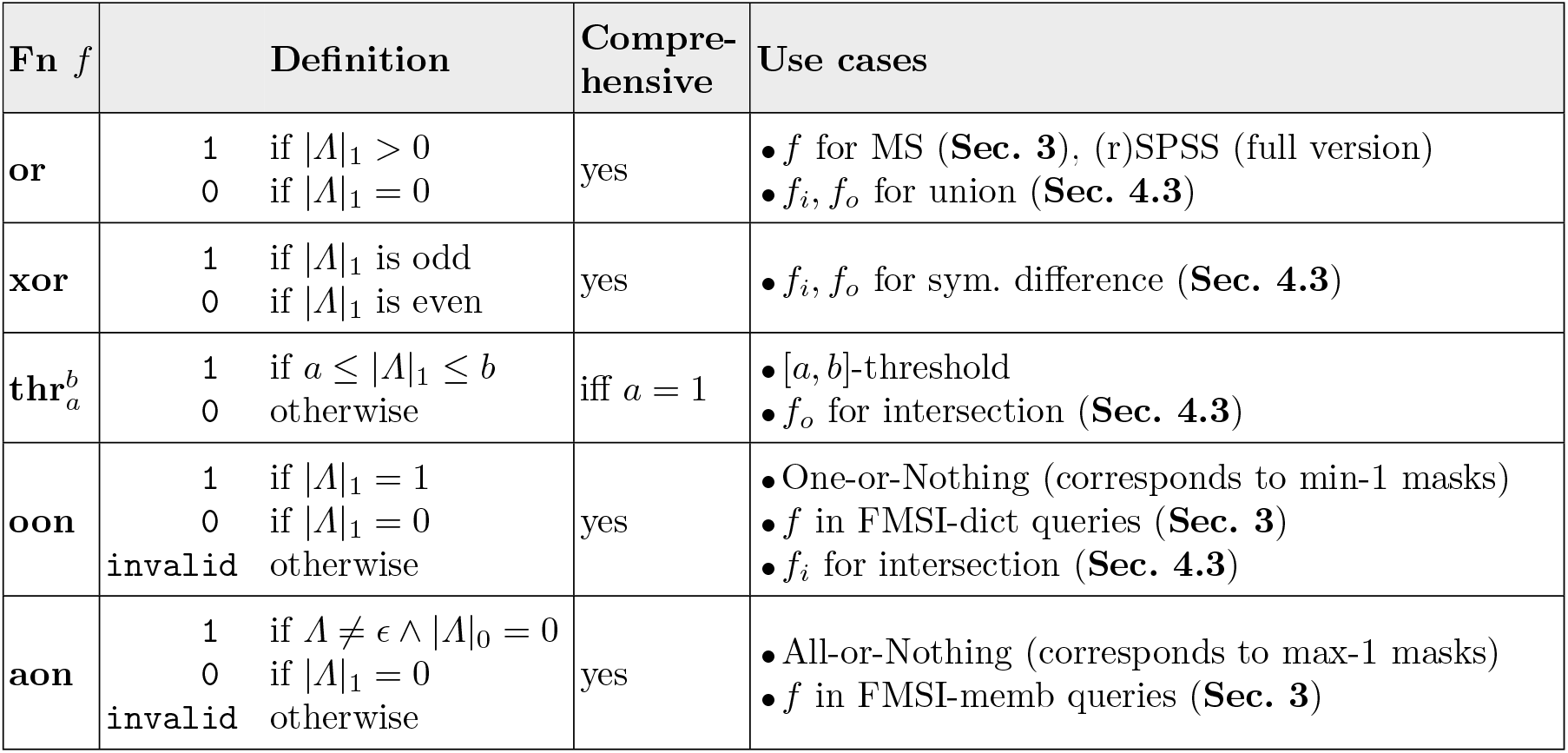
Overview of demasking functions. *Λ*(*f, S, M* ) is abbreviated as *Λ*. Only comprehensive functions can be a target of recasting.

**Figure 1:**
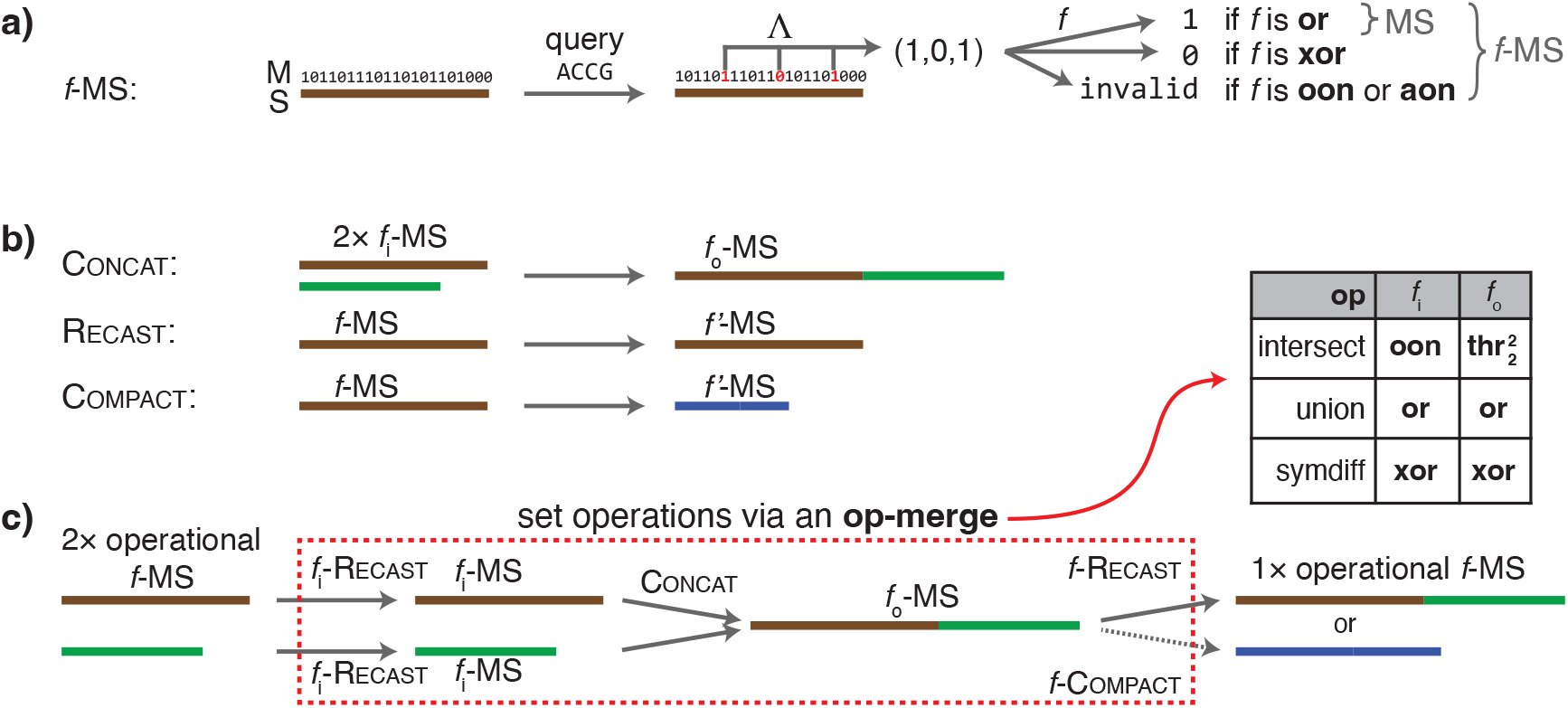
Overview of the *f*-MS framework. **a) The concept of** *f* **-MS**. For a given *k*-mer, the corresponding mask bits *Λ*(*S, M, Q*) are evaluated using a demasking function *f* . **b) Low-level operations**. A *f* → *f* ^*′*^ Recast changes mask under function *f* to another mask under function *f* ^*′*^, while preserving the represented *k*-mer set. Concat merges two superstrings and masks. Both operations may conceptually be performed either on the original *f*-MS or on their associated indexes. **c) Set operations**. An operation **op** is performed by a sequence of Concat and Recast applied to the input *f*-MS, with operation-specific input and output functions (see Tab. 1). The Recast, which serves for keeping the *f*-MS operational for its data structure, may be replaced by compaction with the same target function.

### Example 5.

Consider Example 2 with the set of 3-mers *K* = {`ACG`, `GGG`} and the masked superstring `AcGGgg`. For the query *k*-mer *Q* = `GGG`, the occurrence function gives *Λ*(*S, M, Q*) = (1, 1). The result of demasking *f* (*Λ*(*S, M, Q*)) then depends on the specific function *f* : for **or** it evaluates to 1 and thus `GGG` is represented; however, for **xor** the result would be 0 and `GGG` would not be represented.

### Operational *f*-MS with respect to different applications

Different use cases may require different properties of masks. Since the choice of a demasking function can be also viewed as a restriction of the set of acceptable masks for a given super-string and *k*-mer set, we can choose different demasking function to enforce certain properties of the masks. We call the demasking functions that enforce the desired properties of masks as *operational*. The concept of operational demasking functions plays a central role in the whole *f*-MS framework, see Figure 1c.

As an example, considering the FMSI index [39], for its version for membership queries, only **aon** is operational. This is due to a certain query speed optimization. If this optimization is omitted, all demasking functions are operational. For the dictionary version, only **oon** is operational. For more details, see Section 5.

## 4 Abstract algebraic framework for *k*-mer set operations

The dynamization of a data structure can be conceptually split into two parts – the dynamization of the representation and the dynamization of the data structure internals itself. Here, we give a general approach to dynamize the representation, based on two low-level operations for their manipulation – mask recasting and concatenation, where we assume that the concatenation in the data structure can be done efficiently. In the context of different data structures, these can have very different forms, and we discuss a specific variant for BWT in Section 5.

### 4.1 Concat and Recast: The Two Essential Low-Level Operations

The elementary operations of concatenation and recasting described next are those used to perform set operations, as described in Section 4.3. While concatenation effectively merges two *f*-MS, recasting facilitates adjusting *f*-MS into the desired form, e.g., before executing a set operation or making it operational after performing the operation. See also Figure 1 for conceptual overview of these operations.

#### Concat

For full generality, we define concatenation on *f*-MS as concatenating the underlying superstrings and masks for all possible input and output functions *f* .

##### Definition 6.

*Given f-MS* (*f*_*i*_, *S*_1_, *M*_1_) *and* (*f*_*i*_, *S*_2_, *M*_2_), *we define the* (*f*_*i*_, *f*_*o*_)*-concatenation as the operation taking these two f*_*i*_*-MS and producing the result* (*f*_*o*_, *S*_1_ + *S*_2_, *M*_1_ + *M*_2_).

Note that Definition 6 can be easily extended to more than two input *f*-MS. In the case that all the functions are the same, i.e., *f* = *f*_*i*_ = *f*_*o*_, we call it *f-concatenation* or just *concatenation* if *f* is obvious from the context.

Furthermore, note that while the set of appearing *k*-mers of *S*_1_+*S*_2_ clearly contains the union of appearing *k*-mers of *S*_1_ and of *S*_2_, additional new occurrences of *k*-mers may appear at the boundary of the two superstrings. These newly occurring *k*-mers may not be appearing in any of the superstrings *S*_1_ and *S*_2_. The occurrences of appearing *k*-mers of *S*_1_ + *S*_2_ that overlap both input superstrings are called *boundary occurrences*.

##### Example 7.

Consider **aon**-MS `AcGGgg`, representing 3-mers *K*_1_ = {`ACG`, `GGG` }, and **aon**-MS `Ggg`, representing *K*_2_ = {`GGG` }. Their **aon**-concatenation is AcGGggGgg, with two boundary occurrences of GGG; causing the result to be an invalid **aon**-MS even though the input **aon**-MS were valid. Similarly, boundary occurrences of *k*-mers can change the presence in the result for some functions, illustrating the need of a careful treatment of concatenation.

#### Recast

When performing operations with *f*-MS, recasting is used to change the demasking function *f* without altering the represented *k*-mer set and the underlying superstring; here, we note that recomputing the superstring is the most resource-demanding operation [37].

In many cases, recasting triggers recomputation of the mask, which is however a less expensive operation. We first note that recasting to an arbitrary demasking function may not be possible with a fixed superstring as there is no valid mask, e.g., for a function that requires the represented *k*-mers to have at least two on occurrences. Nevertheless, recasting is possible for a large class of demasking functions, specifically functions with the property that all appearing *k*-mers are encodable using a compatible mask as present or absent, irrespective of the number of their occurrences; we call such functions *comprehensive*:

##### Definition 8.

*A demasking function f is* comprehensive *if for every n >* 0, *there exist x, y* ∈ {0, 1}*n such that f* (*x*) = 0 *and f* (*y*) = 1.

The definition directly implies the possibility of recasting an *f*-MS by changing the function *f* to any comprehensive *f* ^*′*^. This also implies an efficient initialization procedure to create a desired operational *f*-MS for a comprehensive *f* : First, compute an **or**-MS as described in [37] and then recast it to *f* .

##### Lemma 9 (recasting)

*Let* (*f, S, M* ) *be a valid f-MS representing a k-mer set K. Then, for every comprehensive demasking function f* ^*′*^, *there exists a valid mask M* ^*′*^ *such that* (*f* ^*′*^, *S, M* ^*′*^) *represents K*.

*Proof*. Since *f* (*ε*) = 0 for any demasking function *f*, we know that non-appearing *k*-mers are not represented in the *f*-MS (*f, S, M* ) and will correctly be non-represented in the resulting *f* ^*′*^-MS. For any appearing *k*-mer *Q* in *S*, we find all of its *n*_*Q*_ *>* 0 occurrences and then use Definition 8 to set the mask bits of *M* ^*′*^ at these occurrences either to *x* with *f* (*x*) = 0 if *Q* ∉ *K*, or to *y* with *f* (*y*) = 1 otherwise (since *f* is symmetric by Definition 3, the order does not matter).

The proof gives a general algorithm for recasting to any comprehensive function as for each appearing *k*-mer we can check whether it is represented in the *f*-MS of origin, and find the number of on occurrences for it to be represented (or not) in the new *f*-MS. Moreover, for all comprehensive functions mentioned in Table 1, recasting can be done either by maximizing the number of 1s in the mask (**aon**), or by minimizing the number of 1s (all other comprehensive functions in Table 1). Both can be achieved in linear time, namely using a two-pass algorithm for max-ones or a single-pass algorithm for min-ones, as described in [37].

### 4.2 Compact: an Optional Low-Level Operation

After performing a number of set operations, the resulting superstring may be much longer than the size of the *k*-mer set it represents. Then it is desirable to change also the superstring alongside recasting the mask. We call this operation of changing (*f, M, S*) into (*f* ^*′*^, *M* ^*′*^, *S*^*′*^), while preserving the represented set *K, compaction*, and typically require |*S*^*′*^| *<* |*S*| . Lemma 9 implies that if *f* ^*′*^ is comprehensive we can use any superstring of *K* as our *S*^*′*^, and therefore compact can be performed by any algorithm for superstring computation. We give an example of how compaction can be performed on a BWT-based index in Section 5.

### 4.3 *k*-Mer Set Operations via *f*-MS and its Low-Level Operations

#### Union

As implicitly shown in [37], concatenating MS, which are **or**-MS in our notation, acts as union on the represented sets; that is, the resulting represented set is the union of the original represented sets. Specifically, the boundary occurrences of *k*-mers do not affect the result by the definition of **or**. This allows **or**-MS to generalize SPSS representations, since any set of *k*-mers in the SPSS representation can be directly viewed as an **or**-MS by concatenating the individual simplitigs/matchtigs.

We show that **or** is the only comprehensive demasking function that acts as union on the represented sets, up to the interchange of the meaning of 0s and 1s; see Supplementary Materials [38] for details. We further demonstrate this uniqueness even when the concatenated masked superstrings directly correspond to individual matchtigs and, therefore, **or**-MS are the only *f*-MS that generalize SPSS representations.

#### Symmetric difference

Next, we observe that **xor** naturally acts as the symmetric difference set operation, i.e., concatenating two **xor**-MS results in an **xor**-MS representing the symmetric difference of the original sets. Indeed, recall that using **xor** implies that a *k*-mer is represented if and only if there is an odd number of on occurrences of that *k*-mer. Observe that the boundary occurrences of *k*-mers do not affect the resulting represented set as those have zeros in the mask. Thus, if a *k*-mer is present in both sets, it has an even number of on occurrences in total and hence, it is not represented in the result. Likewise, if a *k*-mer belongs to exactly one input set, it has an odd number of on occurrences in this input set and an even number (possibly zero) in the other; thus, it is represented in the result. As any appearing *k*-mer is either boundary or appears in one of the input MS, the result corresponds to the symmetric difference.

#### Intersection

After seeing functions for union and symmetric difference operations, it might seem natural that there exists a comprehensive function *f* such that *f*- concatenation yields intersection. This is however not the case as there exists no comprehensive demasking function acting as intersection, see Supplementary Materials [38]. We can circumvent the non-existence of a single demasking function acting as intersection by using a different function on the output than on the input. To this end, we will need two different demasking functions:

– The **oon** (abbreviation of one-or-nothing) function is a demasking function that returns 1 if there is exactly one 1 in the input, 0 if there are no 1s, and `invalid` if there is more than a single on occurrence of the *k*-mer.
– The 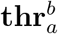 function (an abbreviation of threshold), where 0 *< a ≤ b*, is a demasking function that returns 1 whenever it receives an input of at least *a* ones and at most *b* ones and 0 otherwise. Note that unless *a* = 1, 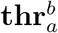 functions are not comprehensive. The corresponding *f*-MS are denoted 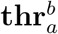 -MS.

We use **oon**-MS to represent the input masked superstrings and treat the result of the concatenation as 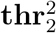 -MS, that is, we apply (**oon**, 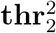)-concatenation. Since the represented *k*-mers are those that have precisely one on occurrence in the original **oon**-MS, they are those with two on occurrences in the result and therefore are correctly recognized by the 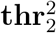 function.

#### Further generalizations and optimization

Other set operations can be performed by chaining intersection, union, and symmetric difference. For example, the asymmetric difference *A −B* can be obtained as *AΔ*(*A∩B*). Such an approach is possible for *any set operation* on *any number of input sets*.

Furthermore, the approach with **oon**-MS for intersection can be utilized to obtain more efficient computation of any symmetric operation for any number of input sets. If we represent each input as **oon**-MS, then after concatenation of all of them, the number of on occurrences counts the number of sets where the *k*-mer appeared and so the symmetric set operation can be recognized with appropriate demasking function. Furthermore, the same such mask and superstring can be used for different operations, e.g., for intersection and union at once, just with different *f* .

Finally, there are more demasking functions that could be used as operational in other use cases. For instance, the **and** function, for which an appearing *k*-mer is represented if all its occurrences are on, could be used to allow on occurrences of non-represented *k*-mers, potentially making the masks of **and**-MS better compressible.

## 5 Example Application: Dynamization of the FMSI Index

The *f*-MS framework may be used for the dynamization of many different data structures. Here, we will demonstrate such a procedure on the FMSI index introduced in our concurrent work [39]. The FMSI index [39] for a *k*-mer set *K* is constructed from its masked superstring (*M, S*) either maximizing the number of ones for membership queries (FMSI-memb), or minimizing them for dictionary queries (FMSI-dict). The FMSI index consists of the Burrows-Wheeler transform (BWT) [8] of *S* with an associated rank data structure, and the *SA-transformed mask M* ^*′*^ which is a bit-vector of the same length as *S*, where *M* ^*′*^[*i*] = *M* [*j*_*i*_ *−*1 mod |*M*| ] where *j*_*i*_ is the starting position of the lexicographically *i*-th suffix of *S*. FMSI can be constructed in linear time through Masked BWT [39], a tailored variant of the classical BWT [8]. FMSI can index a *k*-mer *Q* in *O*(*k*) time by first computing the range of occurrences of *Q* in the suffix-array coordinates. Here, we only consider the most memory-efficient version of FMSI index, which requires 2 + *o*(1) bits of memory per distinct *k*-mer under the spectrum-like property and at most 3 + *o*(1) bits per superstring character in the general case [39]. In addition, we consider the rank data structure also for the SA-transformed mask, which does not asymptotically increase complexity, that is, costs only another *o*(1) bits per superstring character.

### 5.1 Step 1: Extending the FMSI index to *f*-MS

The first step to an *f*-MS dynamization of a data structure is to identify its operational demasking function. For FMSI-memb and FMSI-dict, this is **aon** and **oon**, respectively, but the implementation can be generalized for any demasking function to be operational. Afterwards, we simply implement the evaluation of the operational demasking functions, possibly with some pre- and post-processing which may be specific to the operation and to the internals of the data structure.

#### Efficient membership queries with arbitrary demasking functions

As FMSI-memb retrieves only the mask symbol at an arbitrary occurrence, it requires all the symbols of a *k*-mer to be the same. Thus, the only comprehensive operational *f*-MS is **aon**-MS, which returns 1 if it receives a list of ones, 0 is a list without a one, and `invalid` otherwise (see Table 1). This *f*-MS corresponds to **or**-MS with maximized number of ones.

However, the search in FMSI can be easily extended to return all occurrences of a *k*-mer. In such case, all possible demasking functions are operational. In Lemma 10, we demonstrate that if the function can be evaluated efficiently, *k*-mer presence in FMSI can be determined in constant time after performing backwards search.

##### Lemma 10.

*Consider a query for k-mer Q on an f-MS* (*f, S, M* ) *representing a k-mer set K, such that f can be computed in O*(1) *from the number of* on *and* off *occurrences of Q. Let M* ^*′*^ *be the corresponding SA-transformed mask [39] and assume we know the range* [*i, j*) *of sorted rotations of S starting with a k-mer Q. Then the presence or absence of Q in K can be determined in O*(1) *time*.

*Proof*. From [39, Lemma 1], *M* ^*′*^[*x*] for *x* ∈ [*i, j*) corresponds to the mask symbol of a particular occurrence of *Q*. Therefore, |*Λ*(*S, M, Q*) |_1_ = rank_1_(*M* ^*′*^, *j*) *−*rank_1_(*M* ^*′*^, *i*), which can be computed in *O*(1) time using two rank queries on the mask; here, 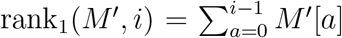 is the number of ones on coordinates 0, …, *i −* 1 in *M* ^*′*^, computed by the rank data structure. Furthermore, |*Λ*(*S, M, Q*) |_0_ = |*Λ*(*S, M, Q*) | *−*|*Λ*(*S, M, Q*) | _1_ = *j −i −*|*Λ*(*S, M, Q*) |_1_. Since *f* is commutative, *f* (*Λ*(*S, M, Q*)) can be evaluated from the two quantities in constant time.

#### Dictionary queries

For dictionary queries, FMSI requires the mask to contain only a single on occurrence of a *k*-mer. This exactly corresponds to **oon**-MS (introduced in Section 4.3, see Table 1). After a recast to **oon**-MS, dictionary queries work in the same way as in FMSI-dict by retrieving the rank in the SA-transformed mask of the first lexicographic occurrence [39].

### 5.2 Step 2: Adding Concat and Recast

#### Recast

Recasting of an *f*-MS can be done directly in SA coordinates without additional memory. For each position of BWT that was not previously visited, we first retrieve the *k*-mer by traversing the inverse LF mapping [39, Alg. 3] and then obtain the range of this *k*-mer. Then from Lemma 9, we find the number of ones needed to make this *k*-mer represented if it was represented in *f*-MS of origin. This takes time *O*(|*S* |+ *k* |*K*^*′*^| ), where *K*^*′*^ are all appearing *k*-mers. Then, we need to update the auxiliary data structures (rank and select), possibly by recomputing them.

#### Concat

Performing the operation on indexes boils down to merging two BWTs using any algorithm for BWT merging, for example [21] which runs in linear time. To merge the SA-transformed masks, we attach the mask symbols to the corresponding characters of BWT, in the same way as for FMSI construction [39, Alg. 1] and perform BWT merge; hence, the existing algorithms for BWT merging can be adjusted in a straightforward way to merge the SA-transformed masks. The BWT merging is then followed by recomputation of the ranks necessary for the functioning of FMSI.

### 5.3 Step 3 (optional): Adding Compact

If an *f*-MS contains too many redundant copies of individual *k*-mers, e.g., if an *f*-MS is obtained by concatenating multiple input *f*-MS, it might be desirable to *compact* it, i.e., reoptimize its support superstring. This can be done by recasting to **oon**-MS, exporting this **oon**-MS to its superstring form by reversing the LF-mapping, removing all segments of support superstring not containing any on occurrence of a *k*-mer, which can be done by single-pass algorithm, and reindexing the result. Although this algorithm requires reindexing of the result, it uses memory linear in the total size of the input superstrings. We leave it as future work to design more efficient algorithms for compaction directly in the FMSI index without the need to export the *f*-MS.

## 6 Implementation and Proof-of-Concept Experiments

To demonstrate feasibility of using *f*-MS for set operations in practice, we developed a prototype implementation of the framework, embedded it into the FMSI *k*-mer-set index [39], and evaluated it using two selected genomes.

### 6.1 Implementation in the FMSI tool

We implemented the *f*-MS framework as an experimental extension of the FMSI program [39] (https://github.com/OndrejSladky/fmsi). Our implementation supports any value of *k* up to 64. In its basic form, FMSI takes an input masked superstring, constructs an FMSI index over it and enables performing membership and dictionary *k*-mer queries. For membership queries, we extended the original FMSI implementation [39, Alg. 3] to reflect the new Lemma 10. Dictionary queries are supported as in the original implementation from [39].

Set operations follow the workflow described in Figure 1. The Concat operation is implemented via FMSI index merging: first, by exporting the individual masked superstrings, then, by concatenating them, and finally by reindexing the newly obtained masked superstring result, with the *f*_*i*_, *f*_*o*_ functions tracked by the user. The Recast operation is currently supported for **or, xor, oon**, and **aon**, and proceeds via three steps: exporting the masked superstring by FMSI, mask re-optimization by KmerCamel [37] (min-1 for **or, xor, oon**, and max-1 for **aon**), and reindexing the new *f*-masked superstring by FMSI. The Compact operation is supported directly in FMSI, and internally runs the global greedy algorithm [37] to approximate shortest superstring.

### 6.2 Experiments using selected genomes

We evaluated our framework using the *C. elegans* and *C. briggsae* genomes, with a primary focus on the memory requirements for processing membership queries. First, we indexed the initial **or**-MS obtained by KmerCamel’s global greedy algorithm [37]. Then, we constructed the indexes for source genomes with *k* = 21 and tested the union, intersection, and symmetric difference operations of the corresponding *k*-mer sets; we note that *k* = 21 is already sufficiently large to capture characteristics of genomes and similar values are typically used for minimizer indexing [31]. Finally, we evaluated the associated quantitative characteristics, including the superstring length of each computed *f*-MS. Additionally, we measured the memory requirements to perform queries on the indexed *f*-MS before and after concatenation, both without and with compaction (measured by GNU time), and also compared FMSI to CBL [28] (commit `328bcc6`, 28 prefix bits). More details are in Supplementary Materials [38].

The starting memory requirements of membership queries of the FMSI and CBL indexes were approximately 2.7 and 83 bits per distinct *k*-mer, respectively, for both of the genomes. After computing the union, this decreased slightly to about 2.5 bits-per-distinct *k*-mer in the union for FMSI, and to about 59 bits for CBL (although still over 20 times more than FMSI). The results for symmetric difference were similar, with FMSI requiring 2.6 bits per distinct *k*-mer and CBL 59 bits per distinct *k*-mer. However, for intersection, which contains only less than 1% of the original *k*-mers, FMSI’s memory consumption increased significantly to 524 bits per distinct *k*-mer, even underperforming CBL whose performance increased to about 420 bits. This is caused by the resulting superstring length being much larger than the represented set, in which case compaction is necessary; indeed, then the memory requirements decreased to 43 bits per *k*-mer (almost 10 times less than CBL). Moreover, in this case, the compaction was relatively cheap thanks to the small size of the resulting k-mer sets compared to the original one. We illustrate this in Supplementary Materials [38].

Overall, this experiment demonstrates that if the percentage of *k*-mers in the result is high, FMSI maintains its superior memory efficiency over CBL even after index merging, without the need for compaction. If the percentage of *k*-mers in the result is very low, compaction needs to be applied in order for FMSI to maintain low memory footprint.

## 7 Conclusion and Outlook

We have proposed *f*-masked superstrings (*f*-MS) as an algebraic data type for *k*-mer sets that allows for seamless execution of set operations. It is primarily based on equipping masked superstrings (MS) from [37] with a demasking function *f*, and we have thoroughly investigated several natural demasking functions, demonstrating that set operations on *k*-mer sets can be carried out simply by masked superstring concatenation or, if indexed, by merging their masked Burrows-Wheeler transform from [39]. This leads to a simple data structure that simultaneously allows for beyond worst-case compressibility, answering exact membership queries, and efficiently performing set operations on the *k*-mer sets, without the costly operation of recomputing the underlying representation. Another major advantage is the versality of our concept as it can in fact be combined with (repetitive) Spectrum Preserving String Sets [10,32,36] instead of (more general) masked superstrings. For instance, our approach can be applied to SPSS-based indexes, such as SSHash – simplitigs of *k*-mers split based on minimizers can be treated as *f*-MS and the same Concat and Recast operations can be performed on those to achieve dynamicity.

The main practical limitation of our work is the current implementation of the index merging, which is very slow and not using the state-of-the-art algorithms for BWT merging [21]. Furthermore, our proof-of-concept experiment is only meant to demonstrate feasibility, and we leave a more thorough evaluation, using various datasets and including a thorough comparison to other tools for set operations, such as CBL [28], to future work.

Our work opens up several research directions for future theoretical investigation. On the algorithmic level, our work relies on efficient algorithms for merging the Burrows-Wheeler transform. Additionally, we seek an algorithm for mask recasting that is not only very space-efficient, but also has faster running time, which can possibly be achieved by using additional data structures for the index, such as the *k*LCP array [34]. Moreover, it is open how to directly perform compaction with *f*-masked superstrings indexed with the masked Burrows-Wheeler transform [39], that does not necessitate to reverse the BWT and export the masked superstring, but rather work locally with the BWT of the superstring and the SA-transformed mask. Furthermore, as our work deals only with set operations, we open the question of performing single insertions and deletions in a more efficient way than performing these through set operations; note that for comprehensive demasking functions, deletions can be handled efficiently by changing the corresponding mask bits.

In conclusion, while the primary contributions of this paper are conceptual, they pave the path towards a space- and time-efficient library for *k*-mer sets that would include all of these features. In the light of advances in efficient superstring approximation algorithms and BWT-based indexing, we believe that the *f*-masked superstring framework is a useful step towards designing appropriate data structures for this library, which is now mainly an engineering challenge.

