## Supplementary Material for "Towards Efficient *k-*Mer Set Operations via Function-Assigned Masked Superstrings" (from another version of this preprint: version 5, 2025-07-07)

#### A Proof of Uniqueness of Union Function

**Theorem 1.** ***or*** *is the only comprehensive demasking function $f$ with $f((1)) = 1$ such that for any two $k$-mer sets $K$ and $K'$ and any of their valid $f$-MS $(f, S, M)$ and $(f, S', M')$, respectively, their concatenation $(f, S + S', M + M')$ is a valid $f$-MS representing the set $K \cup K'$.*

*Proof.* For a contradiction assume there is a comprehensive demasking function $f$ different than **or** that satisfies the above. Consider the smallest $n$ such that $f$ behaves differently than **or** for a length-$n$ input, meaning that there is $x \in \{0, 1\}^n$ not equal to the all-zeros vector such that $f(x) \neq 1$. As $f$ is comprehensive, it cannot happen that $f(x) = 1$ for all $x \in \{0, 1\}^n$, and moreover, $f((1)) = 1$, implying $n > 1$. Fix a $k$-mer $Q$, for simplicity $Q = \mathtt{A}^k$ (although similar approach works for all $k$-mers). We take the first $f$-MS to be the $k$-mer $Q$ with mask being $M_0 = x_0$ and $M_i = 0$ for the remaining $k-1$ positions $i > 0$. The second $f$-MS is $\mathtt{CA}\ldots\mathtt{A}$ where $\mathtt{A}$ appears $n+k-2$ times with the mask being: $M_0 = 0$, $M_i = x_i$ for $i = 1, \ldots, n$, and $M_i = 0$ for $i > n$. At least one of the two represented sets contains the $k$-mer $Q$ as $x \neq 0$ and $n$ is the smallest value for which $f$ differs from **or**. But the resulting $f$-MS is either invalid or does not contain the $k$-mer in the represented set as $f(x) \neq 1$, a contradiction.

**Theorem 2.** ***or*** *is the only comprehensive demasking function $f$ with $f((1)) = 1$ such that for any sequence of $f$-MS, where individual superstrings are matchtigs, the concatenation of all the $f$-MS represents the union of represented $k$-mers.*

*Proof.* It is sufficient to find a construction of matchtigs such that we can construct an arbitrary sequence of ones and zeros at the occurrences of a given $k$-mer and the rest follows similarly as in Theorem 1.

We do this with $k$-mer `CG` and matchtigs `Cg` and `Gc`. Consider the counterexample sequence of occurring ones and zeros from Theorem 1. For every one in the sequence, we add the matchtig `Cg` and for each $m$ consecutive zeros, we add $m + 1$ times the

* Research supported by the French National Research Agency (ANR) under Grant ANR-24-CE45-1226 for the REALL project (KB), by the Czech Science Foundation project 22-22997S (OS, PV), by the ERC-CZ project LL2406 of the Ministry of Education of the Czech Republic (PV), by the Center for Foundations of Modern Computer Science (Charles Univ. project UNCE 24/SCI/008, PV), and by Campus France under PHC BARRANDE 2025 grant n° 52374TC for the EFFIMAS project (KB, PV).

matchtig `Gc`, since at the boundary of two `Gc` matchtigs an OFF occurrence of $k$-mer `CG` appears. At any other boundary, the $k$-mer `CG` does not appear, therefore the construction is correct. The rest of the proof follows a similar argument as in Theorem 1.

Note, however, that the same does not hold if we want to represent simplitigs/SPSS solely. As an individual $k$-mer cannot appear more than once with an ON occurrence, any comprehensive function generalizes SPSS representations if it satisfies that if there is one ON occurrence of a $k$-mer, it returns 1, and if there is none, it returns 0.

### B Non-Existence of a Preserved Comprehensive Function for Intersection.

**Theorem 3.** *There is no comprehensive demasking function $f$ with the property that the result of $f$-concatenation of two $f$-MS always represents the intersection of the originally represented $k$-mer sets.*

*Proof.* Let $f$ be any comprehensive demasking function. Consider MS `A` and `C` with the appropriate mask, each representing a single 1-mer, which from coprehensiveness is always possible. Their concatenation is `AC`. For $k$-mer `A`, the output of occurrence function is the same in `AC` as in `A` and therefore `A` is represented in the concatenation. However, the intersection is empty and thus, $f$ cannot be used to compute the intersection from the concatenation.

Note that the proof cannot be generally extended to all demasking function as there exist non-comprehensive demasking functions acting as the intersection on the represented sets upon concatenation, for instance the constant zero function. However, since the constant zero function always represents the empty set, it is of no use in practice.

### C Details of Proof-of-Concept Experiments

We conducted a simple experiment with genomes of *C. elegans* (`NC_003279.8`, 100M base pairs) and *C. briggsae* (`NC_013489.2`, 108M base pairs), performing the union, intersection, and symmetric difference on their sets of $k$-mers for $k \in \{17, 19, 21, 31\}$. We evaluated the FMSI program [3][1] and compared it with CBL [1][2].

Our proposed pipeline for performing these operations with FMSI, as depicted in Figure 1, consists of five steps: First, we compute a textual representation of the $k$-mer sets interpreted as **or**-MS. In our experiments, this was done using KmerCamel's global greedy algorithm [2]. Second, we recast the mask to the desired demasking function, specifically we keep **or** for union and change to **oon** for intersection and to

[1] Obtained from https://github.com/OndrejSladky/fmsi, version at commit '`778f2f0`'.

[2] Obtained from https://github.com/imartayan/CBL, version at commit '`328bcc6`'. The index was computed on canonical $k$-mers, to handle reverse complements, that is, we ran '`cbl build -c`'. CBL was compiled using '`RUSTFLAGS="-C target-cpu=native" K={kmer-size} PREFIX_BITS=28 \\`
`cargo +nightly build ---release ---examples ---target-dir target.k_{kmer-size}`'.

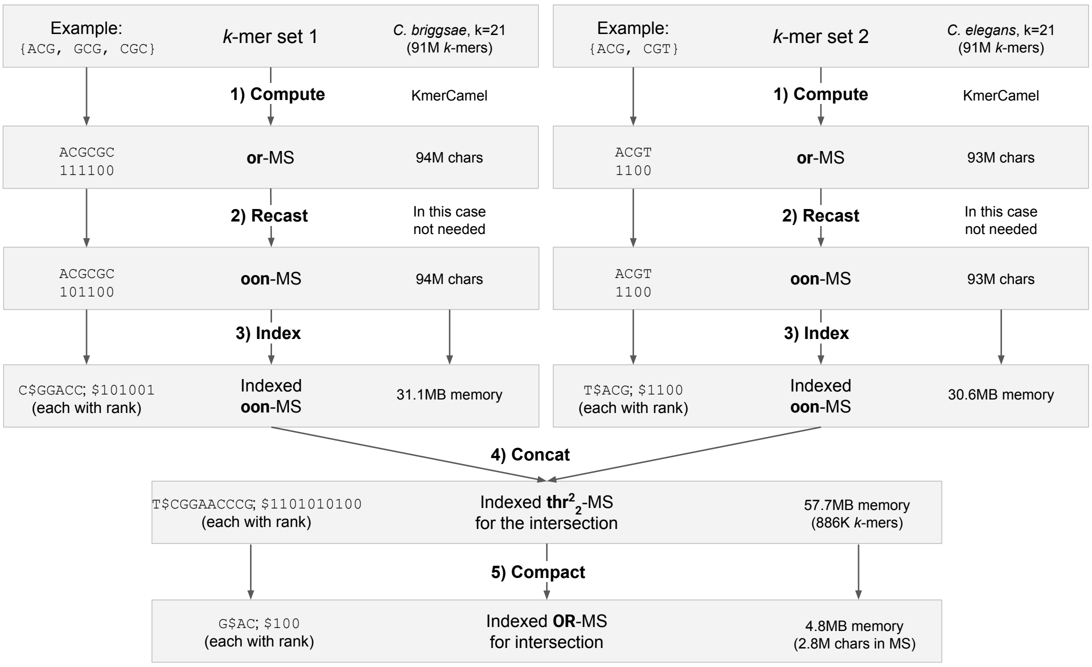


**Figure 1: Illustration of the $k$-mer set operations workflow using $f$-masked superstrings: an example of intersection of two sets using FMSI [3].** The workflow also contains an illustrative example on a set of 3-mers as well as experimental results on *C. briggsae* and *C. elegans* genomes with $k = 21$; namely, we show the number of masked superstring characters after each change, the number of distinct $k$-mers in each set, and for each indexed representation, the memory required to perform queries on the underlying set.

**xor** for symmetric difference. In the case of **or**-MS computed by KmerCamel, mask recasting is actually not needed as the output already minimizes the number of 1s in the mask. Then we index the $f$-MS using FMSI [3] (this can be done even before mask recasting). The last two steps are concatenating the two indexes by index merging and compacting the resulting index if needed. Note that once indexed, we can ask membership queries on the resulting $k$-mer sets.

We evaluated the superstring length of each computed $f$-MS, and the memory requirements to perform queries on the indexed individual and concatenated $f$-MS, both before and after compaction. The results for intersection of the two roundworm genomes for $k = 21$ are depicted in Figure 1. Note also that the fact that compaction significantly reduces the MS length highly depends on the particular use case, namely on the proportion of represented $k$-mers in the result. For union and symmetric difference for the same data, the compaction lead to negligible length reduction.

For results of FMSI with symmetric difference, union, and other values of $k$, see the supplementary repository (https://github.com/OndrejSladky/f-masked-superstrings-supplement).

### References

1. I. Martayan, B. Cazaux, A. Limasset, and C. Marchet: *Conway–bromage–lyndon (cbl): an*

*exact, dynamic representation of k-mer sets.* Bioinformatics, 40(Supplement_1) 06 2024, pp. i48–i57.

2. O. Sladký, P. Veselý, and K. Břinda: *Masked superstrings as a unified framework for textual k-mer set representations.* bioRxiv, 2023.02.01.526717 2023.

3. O. Sladký, P. Veselý, and K. Břinda: *FroM Superstring to Indexing: a space-efficient index for unconstrainedk-mer sets using the masked burrows-wheeler transform (MBWT).* bioRxiv, Nov. 2024.
